# Host cell plasma membrane-derived vesicles efficiently inhibit *in vitro* Influenza A virus infection in a size-dependent manner

**DOI:** 10.64898/2026.05.03.722494

**Authors:** Bushra Qazi, Vaishali Vishwakarma, Vishal Kumar, Garima Pant, Kalyan Mitra, Raj Kamal Tripathi, Sourav Haldar

## Abstract

The influenza virus poses a significant global health threat due to its continuous evolution, immune evasion, and zoonotic spillover. The rise of drug resistance, reduced susceptibility to existing antiviral medications, and the limited effectiveness of annual vaccines underscore the need for new antiviral strategies. To infect, the influenza virus binds to sialic acid (SA)-containing molecules on host cell membranes through hemagglutinin (HA). Blocking this interaction represents a promising antiviral approach. Herein, we report that SA containing plasma membrane-derived vesicles (PMV) efficiently inhibits *in vitro* Influenza A virus (IAV) infection. Using orthogonal methods, we demonstrate that PMV derived from A549, MDCK, and HEK cells competitively bind to H1N1 (WSN) and H3N2 (X-31) IAV strains, block entry and infection in human respiratory epithelial cells in a dose-dependent manner, without causing significant toxicity. When the size of the vesicles was reduced through extrusion, the antiviral activity was enhanced, and this was found to be correlated with a size-dependent increase in hemagglutination inhibition and reduced IAV internalisation. Plasma membrane-derived vesicles may serve as a novel antiviral strategy against influenza virus infections due to their simple production method and conserved SA binding site on HA.

## 1. INTRODUCTION

Influenza A virus (IAV) infection causes respiratory illness, resulting in global morbidity and mortality, and poses a significant economic burden (de Courville et al. 2022). It leads to about a billion cases of seasonal influenza each year, including 3–5 million cases of severe illness and 290,000 to 650,000 deaths from respiratory complications (https://www.who.int/news-room/fact-sheets/detail/influenza-(seasonal)). The 1918 flu pandemic, caused by influenza A virus, infected one-third of the world’s population and is considered the deadliest infection in recorded history (Morens and Fauci 2020; Nogrady 2025). IAV is an enveloped virus from the Orthomyxoviridae family. Its segmented genome encodes eleven proteins, including two surface glycoproteins, hemagglutinin (HA) and neuraminidase (NA). IAV primarily binds to N-acetylneuraminic acid (sialic acid)-containing lipids or proteins on the cell surface of airway and alveolar epithelium via the viral surface glycoprotein hemagglutinin (Sempere Borau and Stertz 2021), (Herold et al. 2015). Following binding, the viral particles are internalized through clathrin-dependent or clathrin-independent endocytosis. En route to the perinuclear region, the viral envelope fuses from within the endosomes with the middle to late endosomal membranes, facilitated by low pH-induced conformational change in HA, to release the viral RNA genome into the host cell cytoplasm (Haldar et al. 2020),(Lakadamyali et al., 2003), (Banerjee et al. 2014). Inside the infected host cells, new viral proteins are synthesized, assembled at the plasma membrane, and progeny viral particles bud out (mediated by NA) to perpetuate the infection cycle. The key steps in the influenza infection cycle are the binding of virus particles to the host cell surface and fusion with the endosomal membranes (Purohit et al. 2025).

Although every stage of the influenza virus infection cycle is druggable (von Itzstein, 2007), (Liu et al., 2023), constant mutations in the viral genome render the Influenza virus resistant or less susceptible to most existing direct-acting antivirals. For example, adamantanes (Influenza ion channel M2 inhibitor) are no longer recommended for the treatment of influenza A Virus (Lampejo, 2020). Likewise, the most effective drugs against Influenza, inhibitors of NA (e.g., Oseltamivir), have become inactive due to resistance (Irwin et al., 2016), (Lampejo 2020). Additionally, antigenic drift and shift can cause new strains to emerge with different tropism and transmissibility. Vaccines are not efficient due to immune escape driven by the accumulation of mutations and must be updated annually (Boni, 2008). Moreover, repeated vaccination against the Influenza virus may lead to a decrease in the vaccine efficacy (Matz & Ellebedy, 2025). Therefore, there is a constant need to develop new drugs, approaches, and treatment options to prevent influenza viral infections and avoid future pandemics (Nogrady 2025).

The first step in the infection cycle of the influenza virus is the binding of viral particles to the cell surface molecules, containing sialic acid, via Influenza HA. Targeting the Sialic acid binding site of HA using naturally occurring and synthetic Sialic acid mimics/ analogs that would prevent the virus attachment to host cell membranes, thus represents a promising antiviral strategy (Carlescu et al., 2009; Costa et al., 2019; Matrosovich & Klenk, 2003). Several natural inhibitors containing SA have been described (Matrosovich & Klenk, 2003). However, the development of small-molecule drugs that inhibit HA-SA interaction has been limited due to low binding affinity (K_d_ approximately at 2−4 mM) between monovalent SA and HA (P. Zhang et al., 2024). To circumvent this issue, a variety of multivalent synthetic polymeric inhibitors such as glycopolymers (polyacrylamides bearing pendant α-sialoside group) (Spaltenstein and Whitesides 1991), sialoside-based linear and dendritic polyglycerol (Bhatia et al. 2017), glycopolymers (Nagao et al. 2019), glycopolypeptides (Ogata et al. 2009), gold nanoparticles and nanogels (Bhatia et al.,2020) have been developed. Recently, a hetero glycopolymer that simultaneously targets NA and HA has been introduced (Parshad et al. 2023). These molecular architectures are designed to introduce multivalent SA residues on suitable polymeric scaffolds to resemble the host cell surface for efficient virus attachment. However, these structures involve multistep synthesis and assembly, a potential bottleneck for clinical development.

In this paper, we have explored the applicability of plasma membrane vesicles from mammalian cells that bear SA-containing lipids and proteins as a novel and facile anti-influenza strategy. Plasma membrane vesicles contain the cell surface molecules, including the receptors for the Influenza virus, without any organelles and mRNA, and can be generated by facile methods with high yield (Le et al., 2021). Using orthogonal approaches, such as, hemagglutination inhibition, virus internalization, viral protein expression by immunofluorescence and western blotting, and virus infectivity assays, our results comprehensively show that plasma membrane vesicles derived from A549, HEK, and MDCK cells efficiently inhibit the H3N2 (X-31) and H1N1 (WSN) Influenza A virus infection in human respiratory cells (A549) primarily by preventing the internalization of virions into cells without exerting any cellular toxicity. This was further confirmed by a significant reduction in viral protein expression. Interestingly, when the size of the vesicles was reduced through extrusion, approximately close to the size of the influenza virus, the antiviral activity was enhanced, and this was found to be correlated with a size-dependent increase in hemagglutination inhibition and reduced IAV internalization. We envisage that our results may lead to a novel anti-influenza strategy.

## 2. RESULTS

### 2.1 Isolation and Characterization of host cell Plasma membrane-derived vesicles (PMV)

PMV were derived from three mammalian cell lines: A549, MDCK, and HEK according to an optimized procedure (see the material and methods sections). PMV were characterized by transmission electron microscopy (TEM) and dynamic light scattering (DLS) analysis. Concentration of the vesicles were determined by BCA assay. The plasma membrane vesicles were found to be largely spherical and polydisperse. The average hydrodynamic diameter (as determined by DLS) was ∼ 300-600 nm, and the zeta potential was around l£10 mV (see Figure 1). We also determined the total Sialic acid levels of these vesicles. The total SA content of these vesicles from different cells was found to be ∼30 nanomoles mg^-1^ of protein.

**Figure 1:**
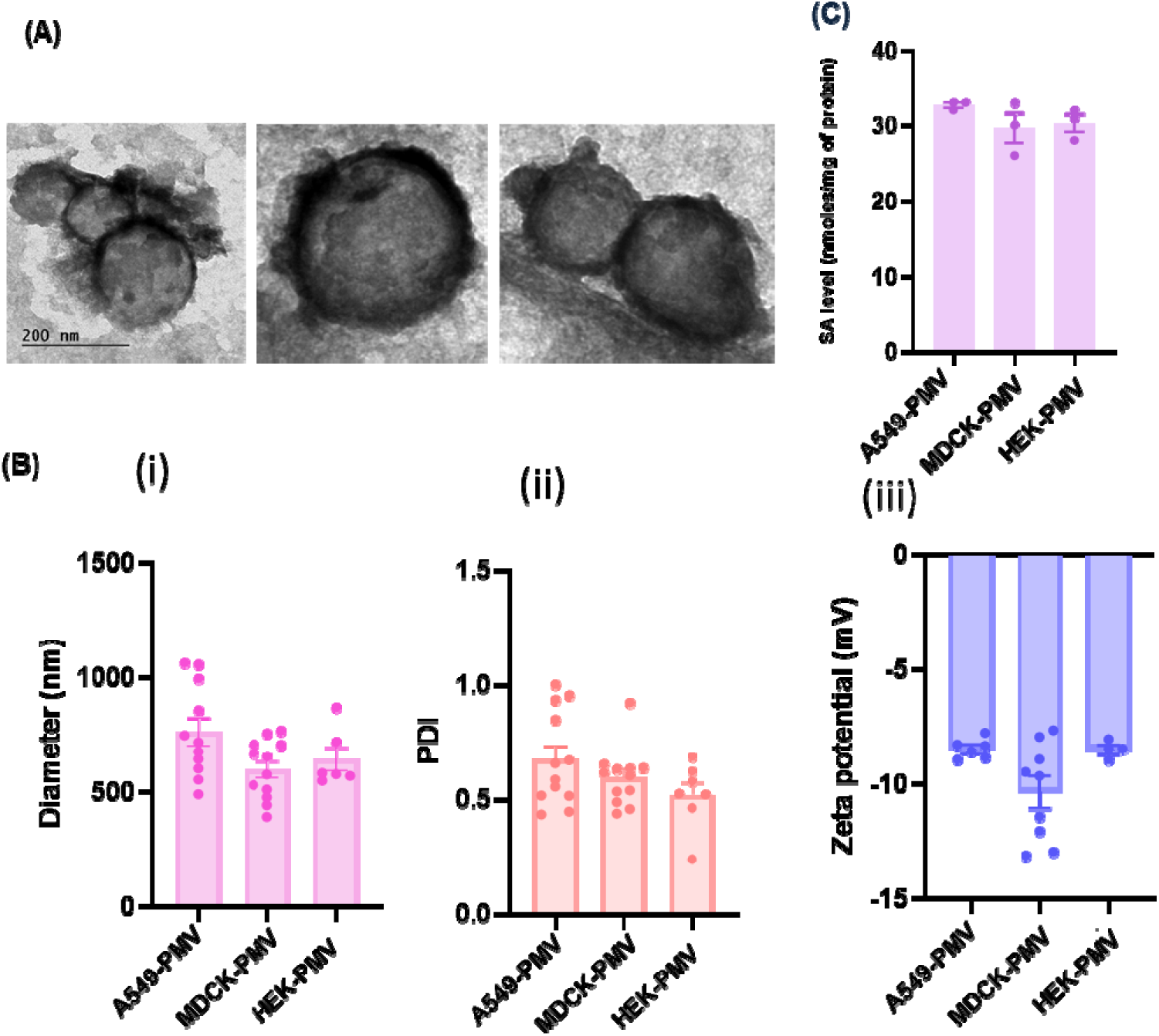
Characterization of PMV. (A) Representative negative stain Transmission Electron Microscopy (TEM) images of extruded plasma membrane vesicles derived from MDCK cells, (B) (i) Diameter, (ii) polydispersity (PDI), and (iii) zeta potential profiles of vesicles derived from A549, MDCK, and HEK cells, (C) Total sialic acid level in PMV. Data represent mean ± SEM of at least three sets.

### 2.2 Influenza A virus binding activity of PMV

To assess the virus-binding activity of plasma membrane-derived vesicles, we carried out a Hemagglutination Inhibition (HI) assay (Pedersen 2014), as shown in Figure 2. HI Assay is a widely used method to evaluate inhibitors of influenza virus binding to its receptors. The assay measures the lowest concentration of an inhibitor that prevents hemagglutination by the influenza virus (Spaltenstein and Whitesides 1991). We performed the HI assay with two different strains of Influenza A virus (X-31/H3N2 and WSN/H1N1) and three types of vesicles (derived from A549, HEK, and MDCK cells). In the absence of PMV, RBCs hemagglutinate by influenza virus and did not precipitate (virus control lane in Figure 2 A, B). However, when viral particles were incubated with plasma membrane vesicles before mixing with RBCs, binding with plasma membrane vesicles prevented the viral particles from agglutinating RBCs, leading to precipitation of RBCs at the bottom of the well, which appeared as a red dot (vesicle lanes in Figure 2A, B). Our HI assay results show that plasma membrane vesicles derived from different cell lines efficiently bind with X-31 and WSN Influenza A virus. The minimum concentration of vesicles for hemagglutination inhibition was found to be ∼ 0.015 mg ml^-1^. Virus binding to plasma membrane vesicles was also confirmed by TEM. Figure 2(C) shows representative negative staining images of virus-bound plasma membrane vesicles.

**Figure 2:**
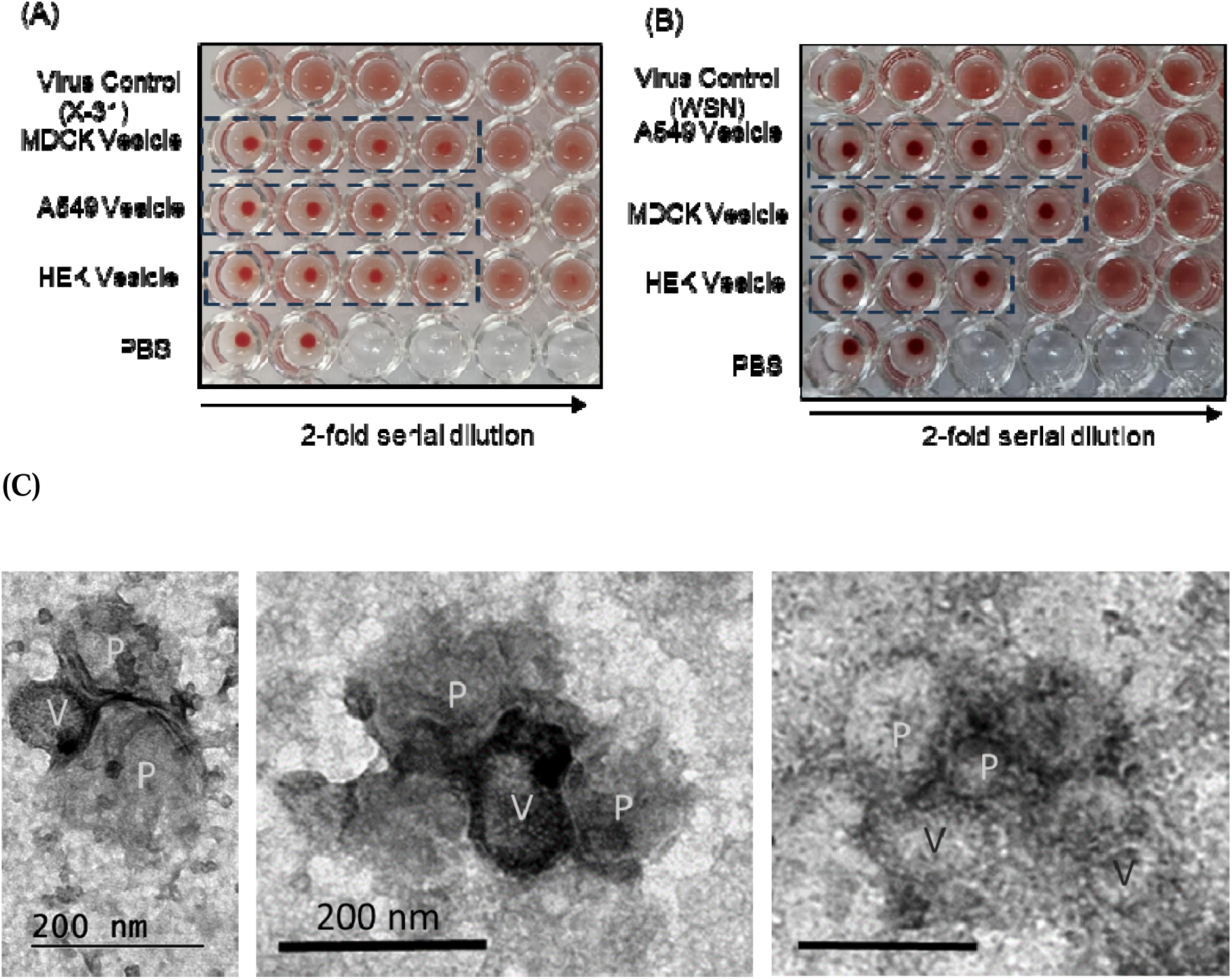
PMV binding to X-31 (A) and WSN (B) Influenza virus by Hemagglutination Inhibition (HI). HI assay was performed with increasing dilutions of PMVs, with only the virus as a positive control and PBS taken as a negative control. The concentration of vesicles was 0.116 mg ml^-1^ for all three vesicles. Wells showing hemagglutination inhibition are marked with dotted lines. See the materials and methods section for details. Influenza A virus (X-31/H_3_N_2_) titer was 256 HA units in 50 µl, and Influenza A virus (WSN/H_1_N_1_) titer was 64 HA units in 50 µl. (C) Representative TEM images showing virus-bound PMV derived from MDCK cells. P = Plasma Membrane Vesicles, V = Influenza A virus (X-31).

### 2.3 Anti-influenza activity of plasma Membrane vesicles

We aimed to leverage the affinity of the Influenza A virus for sialic acid-containing proteins and lipids on the plasma membrane. We hypothesize that vesicles derived from the plasma membranes, which present these viral receptors, can act as decoys to sequester virus particles. Therefore, binding to these vesicles instead of the host cell surface prevents the virus from initiating infection. We tested this hypothesis using two permissive mammalian cell lines, A549 and MDCK, which are commonly used as an influenza virus infection model system. We wanted to ensure that the plasma membrane vesicles are not cytotoxic. Our results show that the plasma membrane vesicles did not exert significant cytotoxic effects in A549, MDCK and RAW 264.7 cells at vesicle concentrations up to approximately 0.12 mg ml^-1^ (Figure S1). To evaluate their antiviral effect, Influenza virus was incubated with varying amounts of PMV derived from A549, MDCK and HEK cells. Next, A549 (or MDCK) cells were treated with infection media containing X-31 (H3N2) or WSN (H1N1) influenza A virus and varying amounts of the PMVs. Light microscopic images show that vesicles significantly reduce the virus-induced cytopathic effect (CPE) with an increase in the vesicle concentration, as shown in Figure 3(A). To further evaluate the anti-viral effect of PMV, we collected media from the infected dish after 48 hours of infection and proceeded with the hemagglutination assay (HA). It is a rapid and reliable test to estimate the virus yield as HA titer. Figure 3(B) shows HA titer decreased in the presence of PMV, suggesting inhibition of IAV infection in A549 cells by PMV.

**Figure 3:**
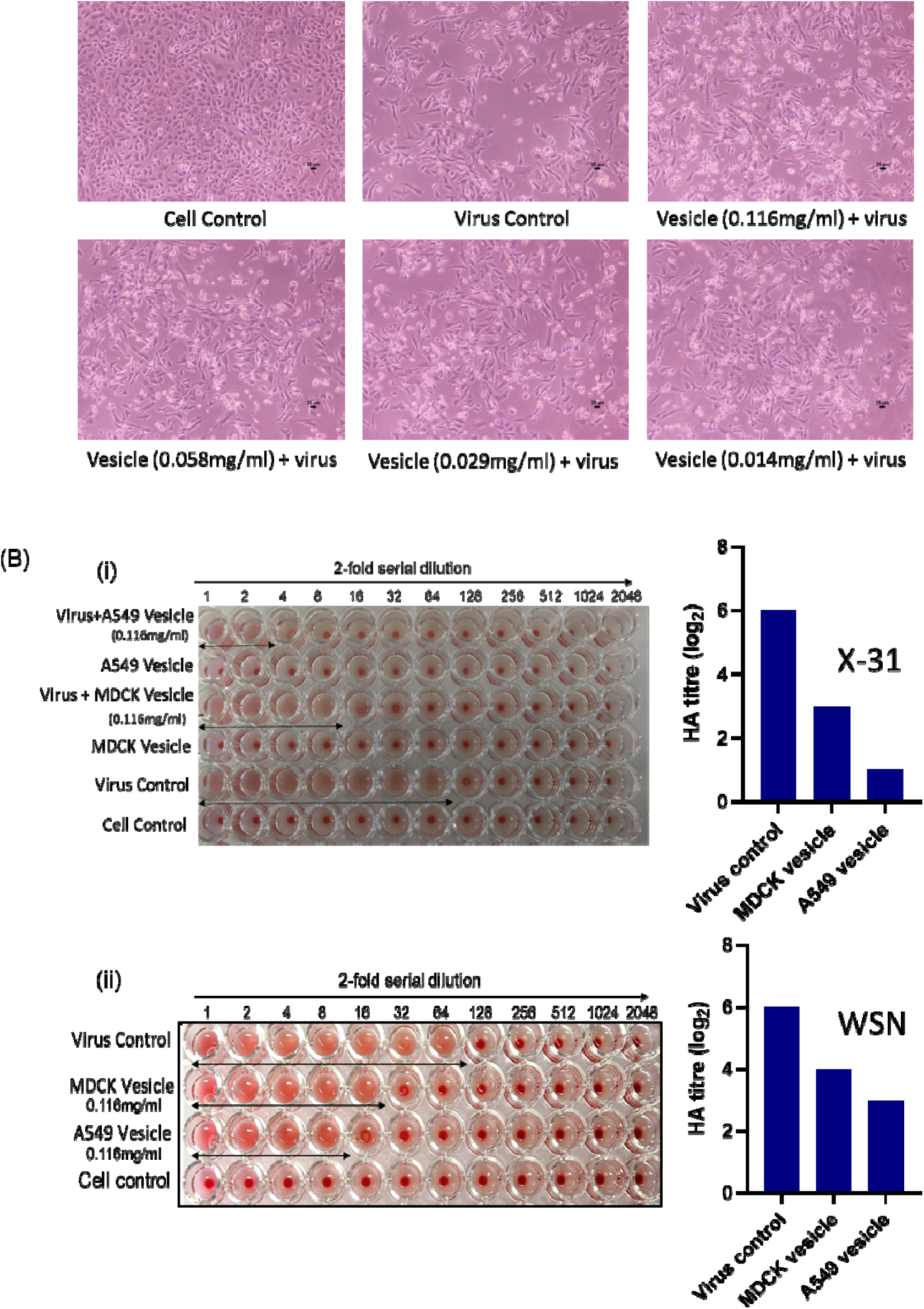
*In-vitro* inhibitory effect of PMV against Influenza A virus infection. (A) Light microscopy observation of reduction in virus-induced cytopathic effect in the presence of A549 vesicle in A549 cells after 48 hours of infection. A549 cells were infected with X-31 (H_3_N_2_) influenza A virus in the absence and presence of 0.116 mg ml^-1^ PMVs. The scale bar is 25μm. (B) Reduction in HA titer after 48 hours post-infection with X-31(i) and WSN (ii) in the presence of 0.116 mg ml^-1^ PMVs derived from A549 and MDCK cells.

To quantify the antiviral effect of PMV, we estimated the reduction in infection in terms of protection from the virus-induced cytopathic effect (CPE). A549 (or MDCK) cells were infected with either X-31 (H3N2) or WSN (H1N1) influenza A virus in the presence of varying concentrations of PMVs. After 48 hours, cell viability was assessed using the MTT assay to evaluate the antiviral efficacy of PMV (Figure 4A, B). The results show a significant reduction in viral infection in the presence of vesicles, as shown by the reduction in CPE in a dose-dependent manner (Figure 4C, D). Our results show that PMVs derived from A549, MDCK, and HEK cells efficiently inhibit X-31 and WSN influenza virus infection in A549 cells. Similar antiviral effects were observed when the infection was carried out in MDCK cells (Figure S2). Interestingly, subsequent addition of PMV after infection (in the presence of PMV) did not have any significant effect on the antiviral activity of the PMV.

**Figure 4:**
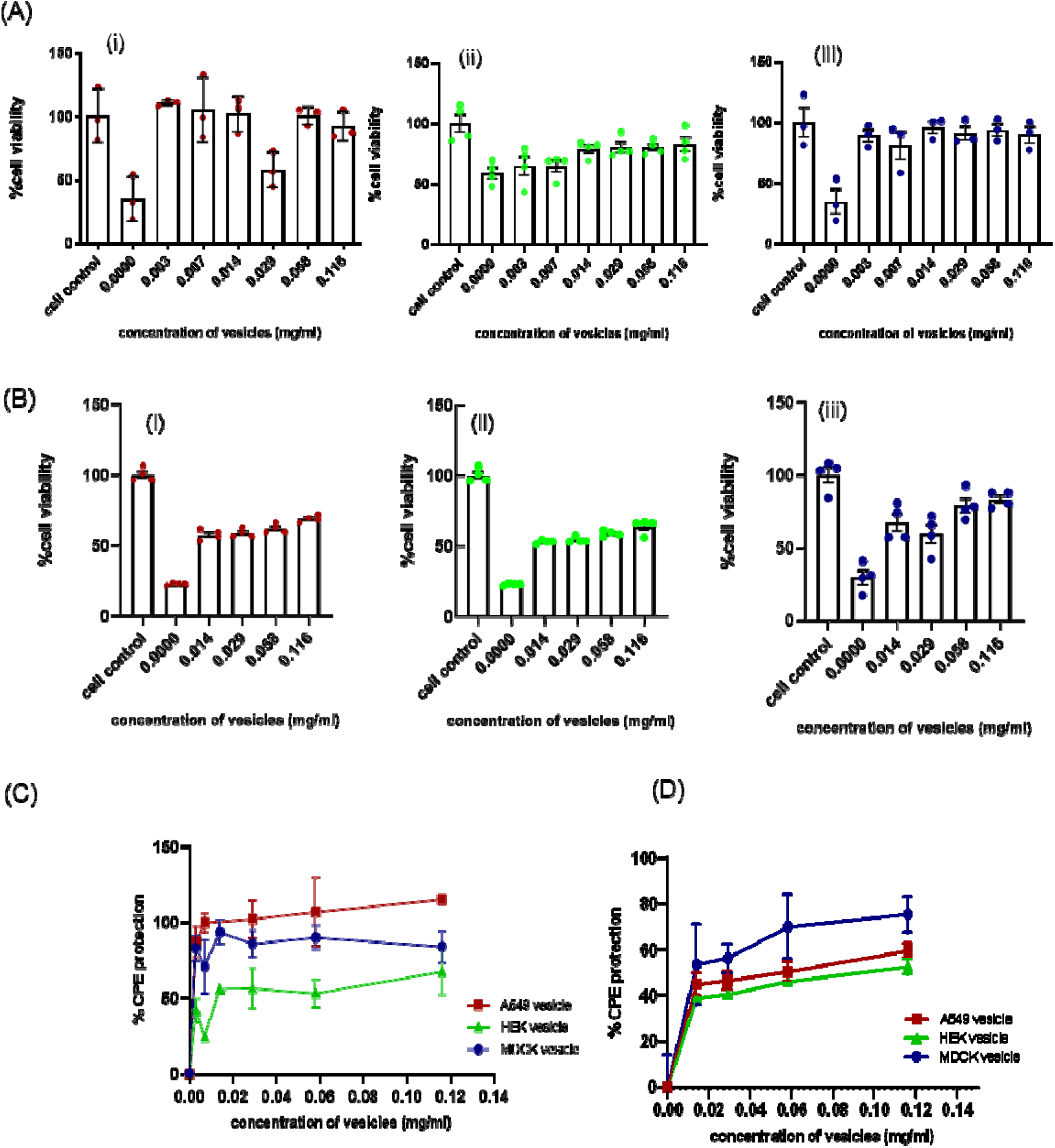
Quantification of *in-vitro* anti-influenza activity of PMV in A549 cells. (A) Protection from X-31 virus-induced CPE (increase in cell viability) by PMV derived from (i) A549, (ii) HEK, and (iii) MDCK cells. (B) Protection from WSN virus-induced CPE by PMV derived from (i) A549, (ii) HEK, and (iii) MDCK cells. A549 cells were infected with X-31 or WSN influenza virus in the presence of varying concentrations of the PMV (derived from A549, MDCK, and HEK). After 48 hours of infection, cell viability was measured by MTT assay. Cell viability was normalized to the control and is reported as the mean ± SEM across at least 3 sets. Dose-dependent reduction in (C) X-31 and (D) WSN infection in A549 cells, as measured by protection from CPE, as a function of PMV concentration. See the materials and methods section for details and estimation of % CPE protection.

The antiviral activity of PMV was further confirmed by a decrease in the expression of the Influenza viral matrix protein M1, as demonstrated by western blot analysis conducted with and without PMV treatment. Figure 5 shows a dose-dependent reduction in the influenza M1 level in the presence of PMV compared to the virus control in A549 cells.

**Figure 5:**
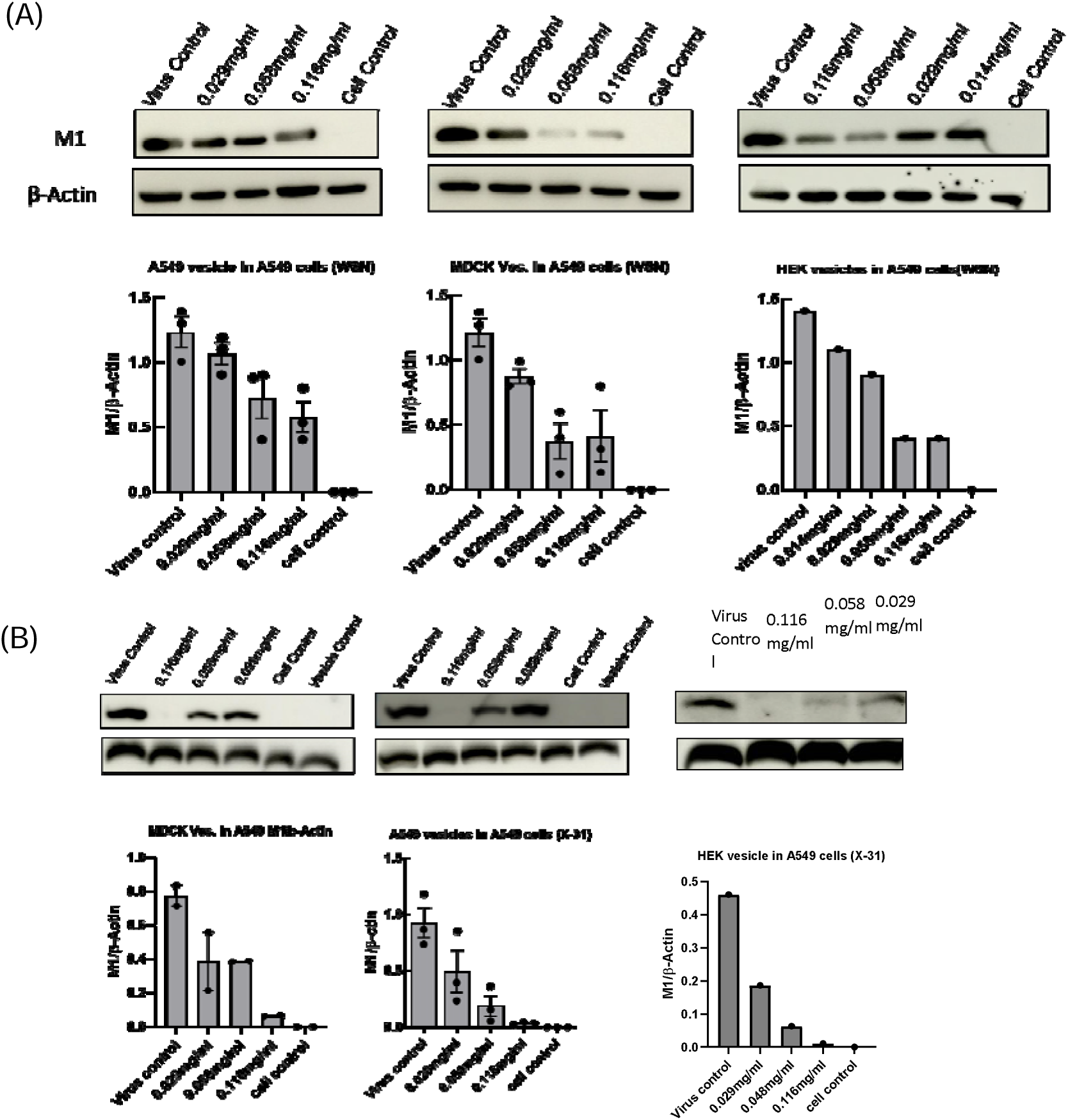
Dose-dependent reduction in influenza A Virus M1 protein levels by PMV in virus-infected cells. Representative western blots with quantification of M1 levels after 48 hours of infection in (A) WSN-infected A549 cells, (B) X-31-infected A549 cells. β-actin was taken as a loading control. Data represent mean ± SEM of three biological replicates in most cases. For the WSN (H1N1) blot, a rabbit polyclonal anti-M1 antibody (1:500) (GTX-127356) and goat anti-rabbit pAb IgG HRP (ab6721) were used. For X-31 (H3N2) blot, a monoclonal anti-M1 antibody (1:1000) (ab22396) and a rabbit anti-mouse IgG H&L [HRP] (ab6728) were used. See the method section for details. Quantification data represent mean ± SEM.

In addition, we performed immunofluorescence staining for M1 protein in A549 cells to evaluate the inhibition of influenza A virus (X-31/H_3_N_2_) infection by PMV derived from A549, MDCK, and HEK cells. A549 cells were infected with the virus in the presence of 0.12 mg ml^-1^ PMV. 15 hours post-infection, immunostaining revealed a significant reduction in infection by all three kinds of PMV. The proportion of infected cells (M1 positive) was quantified and is shown in Figure 6(B). Similar reduction in infection was observed with the (X-31/H_3_N_2_) nucleoprotein (NP) staining (Figure S3).

**Figure 6:**
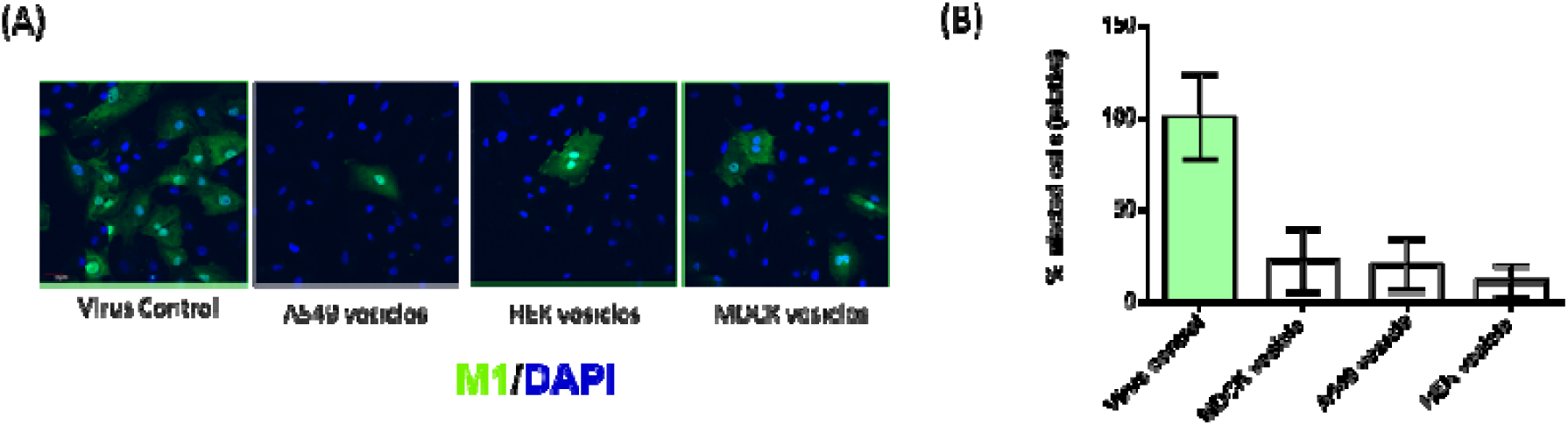
PMV-induced reduction in Influenza A viral (X-31/H_3_N_2_) infection in A549 cells. Infected cells were identified by confocal imaging of the Influenza M1 protein (green) while cells were marked by DAPI (blue) (A). Representative confocal microscopic images showing a decrease in M1 viral protein expression; **(B)** Percentage of infected cells in the presence of PMVs based on M1 staining. (Data represent mean ± SEM of different fields).

### 2.4 PMV inhibits internalization of IAV into cells

According to our working hypothesis, PMV inhibits influenza A virus binding to host cells, thereby preventing viral entry by endocytosis. To test this, we focused on the early binding and entry steps of the viral life cycle. We pre-incubated virus (X-31 (H3N2) or WSN (H1N1) strains) with different concentrations of PMVs at 4°C for 1 hour. A549 cells were then inoculated with the PMV-virus mixture. After 45 minutes of infection at 37°C, 5% CO_2_, we harvested the cells and estimated the intracellular M1 level. Under these conditions, M1 protein only from incoming viral particles would be detected (Eierhoff et al., 2009, 2010). We found a significant reduction in M1 levels when the cells were infected in the presence of PMV compared to untreated controls, suggesting that fewer viruses had successfully entered the cells (Figure 7A, B). Similar dose-dependent entry inhibition of X-31 (H3N2) by PMV derived from A549 cells was also observed (Figure S4). This was further supported by confocal microscopy images, which show fewer viral particles (stained with M1 antibody) inside the cells when the infected in the presence of PMV (Figure 7C). These findings suggest that PMV prevent the virus from effectively binding and entering the host cells. The punctate staining pattern of M1 (after 45 min of infection) compared to diffused staining (after 15 hours of infection, see Figure 6 (A)) indicates that we did not detect any de novo M1 expression in this condition (Banerjee et al. 2014).

**Figure 7:**
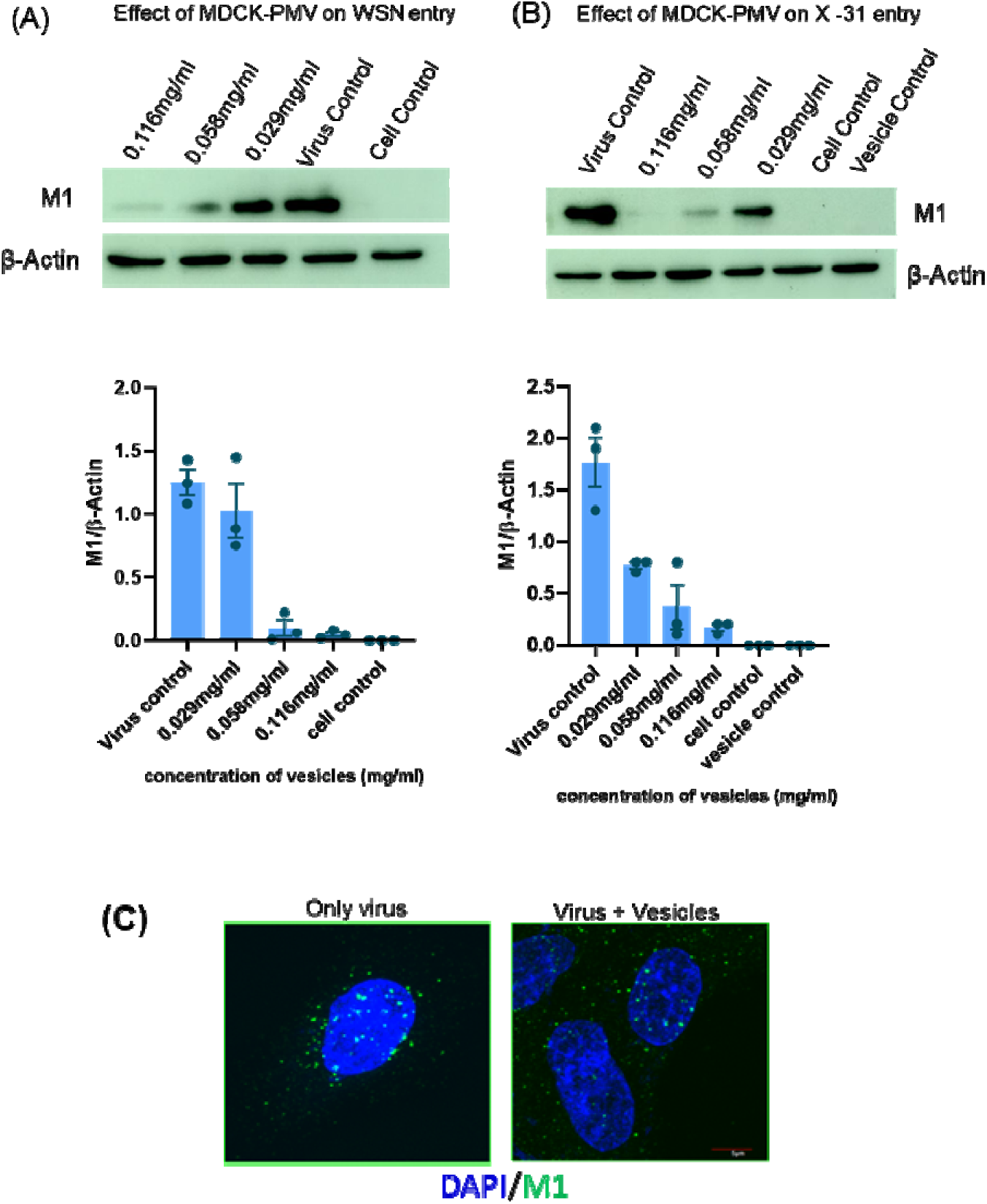
PMV restricts viral entry: dose-dependent reduction in influenza A M1 level (45 minutes post-infection) in A549 cells in the presence of PMV derived from MDCK cells, shown by western blotting. Representative western blot and corresponding quantification of (**A)** Influenza A/ WSN/ H_1_N_1_ and **(B)** Influenza A/ X-31/ H_3_N_2_ M1 levels in A549 cells in the presence of PMV. β-actin was taken as a loading control. Quantification data represent mean ± SEM of three independent experiments. **(C)** Distribution of M1 (X-31 influenza A viral particles) in A549 cells in the presence and absence of MDCK vesicles. Images are maximum-intensity projections along the Z direction.

### 2.5 PMV demonstrates size-dependent antiviral efficacy

To check whether the virus-binding and inhibitory efficacy were dependent on the size of the plasma membrane vesicles, we generated PMV of two different sizes by extruding the crude vesicles derived from the cells. First, we extruded the crude vesicles through 400 nm pore-size filters, then through 100 nm pore-size filters. Figure 8(A) shows the size distribution profiles of crude, 400 nm-, and 100 nm filter-extruded vesicles. We compared the hemagglutination inhibition by crude, 400 nm-filter-extruded, and 100 nm-filter-extruded MDCK vesicles using X-31 and WSN influenza A virus. Our results show that hemagglutination inhibition was highly dependent on PMV size. A clear inverse correlation was observed between vesicle size and binding efficiency; smaller (more uniform) PMVs exhibited significantly improved binding capacity, as evidenced by the decreasing HI titer. Concomitantly, we found that inhibition of viral entry (in terms of intercellular M1 level after 45 minutes of infection) depended on the size of the vesicles (Figure 8C). We therefore expected that the size-dependent binding and entry inhibition would be reflected in the viral infectivity assay. Figure 8 (D) shows size-dependent inhibition of IAV (WSN) infection in A549 cells (CPE reduction) by MDCK vesicles. 100 nm filter-extruded MDCK cell-derived PMV yielded the highest antiviral activity in terms of reduction in cytopathic effect. A549 and HEK vesicles showed similar size-dependent inhibition of IAV endocytosis (Figure S5).

**Figure 8:**
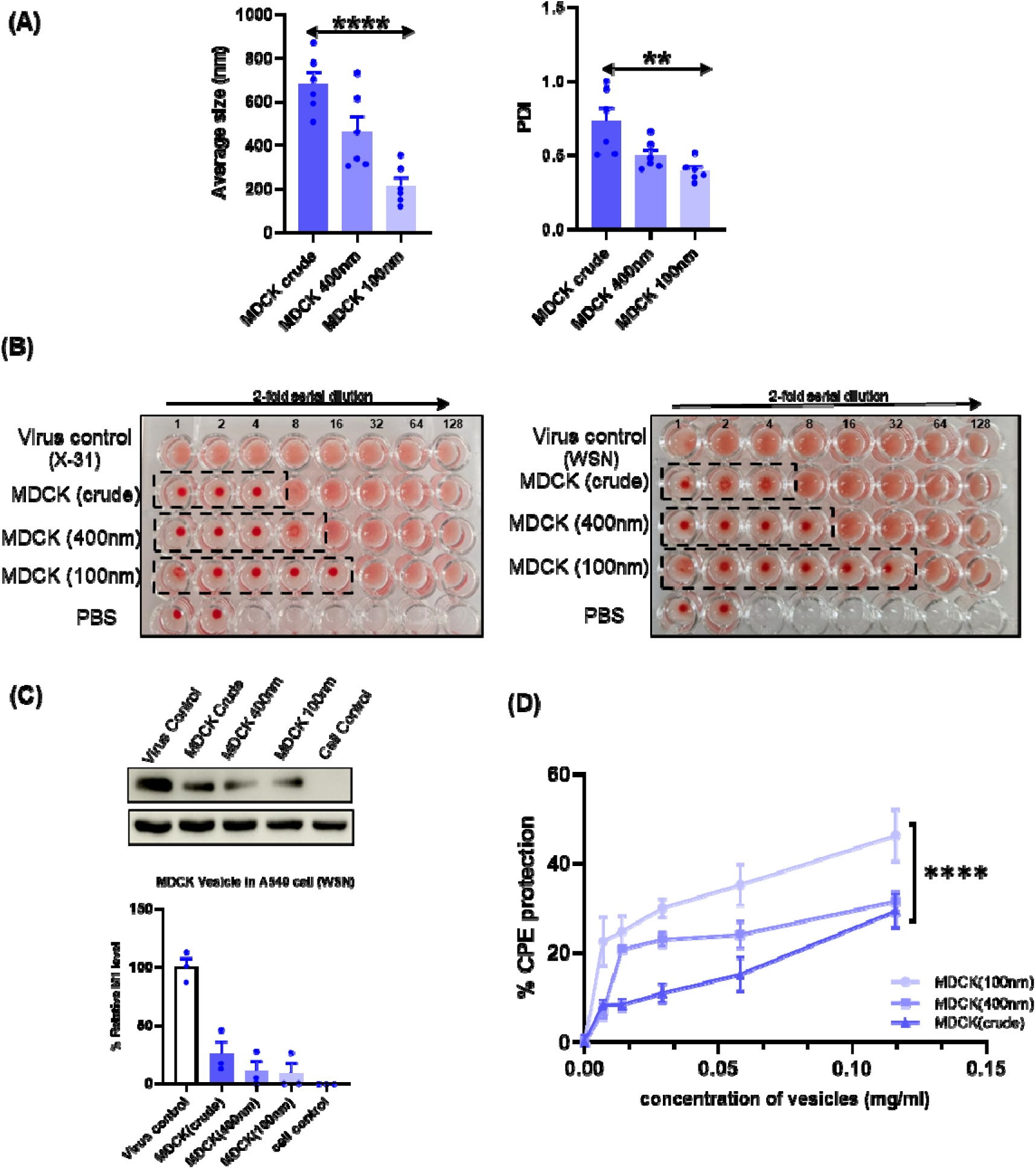
PMV demonstrates size-dependent antiviral efficacy against Influenza A virus infection. **(A)** The average size and polydispersity of the extruded MDCK vesicles. **(B)** The HI assay was conducted with increasing dilutions of MDCK vesicles: PMV (crude), PMV (400nm-filter-extruded), and PMV (100nm-filter-extruded), with only the virus (X-31 and WSN) as a positive control, and PBS as a negative control. **(C)** Influenza A/WSN/ H1N1 M1 protein expression levels in A549 cells in the presence of PMV of varying sizes, as determined by western blotting (45 minutes post-infection). β-actin was used as a loading control. **(D)** Reduction in H1N1 (WSN) infection in A549 cells. Protection from CPE as a function of vesicle concentration by crude,400nm-filter extruded, and 100nm-filter extruded PMV from MDCK cells. Data represent the mean ± SEM of at least three independent experiments (****P<0.0001).

## 3. Discussion

To initiate infection, the influenza virus binds to sialic acid (SA)-containing glycoproteins and glycolipids on the host cell plasma membrane through HA. Although the Influenza genome constantly mutates, the amino acid residues of HA involved in binding with sialic acid are conserved across all subtypes of influenza, throughout antigenic variation (Skehel and Wiley 2000). One approach to abrogate influenza viral infection is to inhibit the virus from binding to cell membranes by competitive binding to viral proteins on the virion surface with decoy receptors. Using a gamut of methods, we demonstrate that SA-bearing plasma membrane vesicles from MDCK, HEK, and A549 cells efficiently bind to and inhibit influenza A virus infection in human respiratory cells (Figure 9). Plasma membrane vesicles are likely to harbour other cell surface receptors (Sempere Borau and Stertz 2021) that may be involved in binding of IAV. Production of plasma membrane vesicles (or Chemical EVs) can be easily scaled up using bioreactors, and are being explored for various therapeutic applications (Alter et al. 2023) (Ingato et al. 2018; Ng et al. 2022). Moreover, PMV-bound viruses can be degraded by macrophages or cleaned off by the reticulo-endothelial system (Longmire et al. 2008); (Bhatia et al. 2016). Virus-bound PMV may also be removed by mucociliary clearance (X. Zhang et al., 2025).

**Figure 9:**
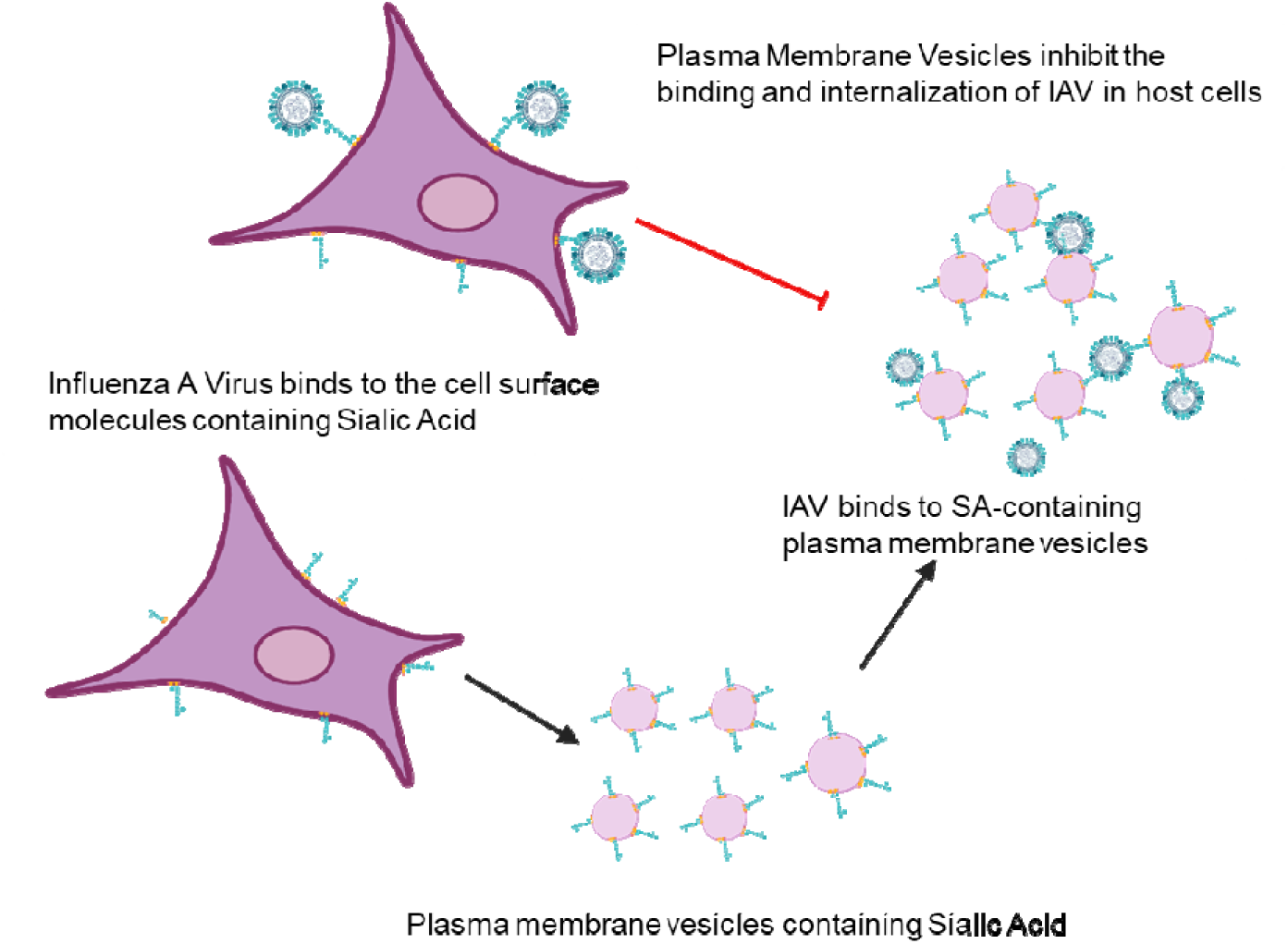
Schematic illustration showing how plasma membrane–derived vesicles (PMV) inhibit influenza virus infection in cells by functioning as sialic acid-rich decoys that prevent virus binding to host cell receptors.

The development of HA-targeting small molecule drugs to inhibit HA-SA interaction has been limited due to low binding affinity (K_d_ approximately at 2−4 mM) between monovalent SA and HA (P. Zhang et al., 2024). In this context, there are several advantages of vesicle-based systems over soluble inhibitors. Vesicle-based systems not only increase the 2D concentration of SA (Bai et al. 2025), but also provide steric shielding and multivalency (Bhatia et al. 2016). Although the interaction of a single HA binding site with a single SA moiety (in solution) is weak, the interaction among multiple HA (on virion surface) and SA on vesicle surface enhances the affinity of virus for SA-containing decoy receptors (Sigal et al. 1996).

In this study, we present a comprehensive evaluation of the *in vitro* anti-influenza activity of plasma membrane vesicles (PMVs) from A549, HEK, and MDCK cells against two representative subtypes of influenza A virus (IAV): A/WSN (H1N1) and A/X-31 (H3N2). Our results indicate that plasma membrane vesicles primarily act as entry inhibitors, which prevent binding and subsequent internalization of virus particles. It is also plausible that internalized PMV-bound virus particles could not fuse with the limiting membranes of the endosomes, and the viral genome is not released in the cytoplasm of the host cells. The binding, entry inhibition, and antiviral activity of PMV were found to be dependent on their size. Smaller vesicles are found to be more efficient in bringing out antiviral activity. This could be since smaller vesicles have a larger surface area, which enhances virus-binding. The ameliorated binding could also be due to the high curvature of the smaller vesicles that influences the orientation of the sialic acid residues on the vesicle surface. It was shown that spherical inhibitors greater than the size of the virus do not benefit from the polyvalent enhancement effect, because there is no increase in the contact area between virus and the nanoparticle, which makes virus-sized nanoparticles/ vehicles the most efficient inhibitors (Bhatia et al. 2016). Extrusion through a 100 nm pore diameter filter generated PMV in the size range of influenza viral particles (100-200 nm), thereby rendering them more efficient entry inhibitors. The higher curvature of smaller PMV may enhance the virus–SA binding by reducing the diffusion of SA-containing proteins or lipids and increasing the probability of attachment (Cail and Drubin 2023).

Although the current study presents promising *in vitro* results, *in vivo* validation of PMV efficacy remains a crucial next step. Future investigations will focus on tracing the biodistribution, pharmacokinetics, and therapeutic efficacy of PMV in animal models of influenza infection. Influenza viral infection is not only a global health hazard but also an economic burden (Putri et al. 2018). The frequent antigenic drift and shift in influenza A virus (IAV), coupled with the increasing emergence of drug-resistant strains, further exacerbate the challenge of effective disease management, underscoring the need to develop alternative, sustainable antiviral strategies beyond conventional small-molecule inhibitors, prone to drug resistance. In this context, we show that host cell plasma membrane vesicles have robust *in vitro* antiviral activity against Influenza A virus, and we propose that sialic acid-containing plasma membrane vesicles can be delivered intranasally for the treatment of Influenza. Keeping in mind the various applications of cellular vesicles (Alter et al. 2023) (Ingato et al. 2018; Ng et al. 2022), We envisage that our antiviral strategy against the Influenza A virus using plasma membrane-derived vesicles can lead to new therapeutics for tackling pandemic influenza.

## 4. Conclusion

In this study, we demonstrated that plasma membrane-derived vesicles (PMV) isolated from human cell lines can effectively block the entry of the influenza A virus into human respiratory cells, thereby inhibiting infection. Importantly, we found that the antiviral effectiveness of PMV depends on their size. We believe that plasma membrane-derived vesicles have the potential to lead to the development of new antiviral therapies.

## 5. Experimental methods

### 5.1 Cell lines, Viruses, and Antibodies

Madin-Darby canine kidney (MDCK), human lung adenocarcinoma epithelial (A549), human embryonic kidney (HEK) 293T cells, and RAW 264.7 cells were obtained from ATCC. All cells were maintained in DMEM (GIBCO) with 10 % FBS (HIMEDIA) containing 100 μg/ml penicillin and 100 μg/ml streptomycin (HIMEDIA). RAW 264.7 cells (macrophage) were maintained in RPMI media (GIBCO) with 5% FBS (HIMEDIA) containing 100 μg/ml penicillin and 100 μg/ml streptomycin (HIMEDIA). Cells were grown in a humidified atmosphere (95% humidity) at 37 °C and 5% CO_2_. Trypsin was obtained from HIMEDIA, and TPCK-Trypsin from Thermo Scientific. Influenza A virus (strain X-31, A/Aichi/68, H3N2) was a kind gift from Dr. Joshua Zimmerberg at NICHD/ NIH, Bethesda; Influenza A virus (strain WSN, H1N1) was a kind gift from Dr. Arindam Mondol, IIT Kharagpur. Mouse monoclonal Anti M1 antibody (ab22396), Rabbit anti-mouse IgG H&L [HRP] (ab6728), goat anti-rabbit pAb IgG HRP (ab6721) were obtained from Abcam. A rabbit polyclonal anti-M1 antibody (GTX-127356) was obtained from Gene Tex. and HRP-conjugated anti-β-actin (A3854) was obtained from SIGMA.

### 5.2 Preparation of Plasma Membrane-derived Vesicles (PMV)

PMV were prepared according to the reported methods with modifications wherever required (Alter et al., 2023; Gray et al., 2015; Sezgin et al., 2012. Briefly, cells (MDCK, A549, and HEK) were grown in DMEM (GIBCO) with 10% FBS (HIMEDIA) and 100 µg ml^-1^ penicillin-streptomycin (HIMEDIA). After they attained 70%-80% confluency, vesiculation was induced. Cells were washed gently with buffer A (2mM CaCl_2_, 10mM HEPES, 15mM NaCl, pH 7.4) two times and then with freshly prepared buffer B (3mg DTT (SRL), 20.3µl of 37% formaldehyde in 10ml of buffer A), followed by an incubation at 37°C for 90-120 minutes at 100 rpm (Kϋhner shaker) in the appropriate quantity of the buffer B. The buffer containing vesicles were collected and centrifuged at 500 RCF for 5 minutes to pellet cellular debris. After separating the cellular debris, the supernatant was centrifuged at 21,000 RCF, for 1 hour at 4°C, to concentrate the vesicles. The pellet was resuspended in 1XPBS and centrifuged at 21,000 RCF for 1 hour at 4°C. Finally, the pellet was resuspended in a minimum volume of PBS and stored at 4°C. The concentration of PMV was determined by bicinchoninic acid assay using a kit (Thermo Scientific, USA).

### 5.3 Characterization of PMV

Physicochemical characterization of PMVs was done by diluting the vesicles in 1X PBS and measured in Zetasizer. Mean hydrodynamic diameter of the vesicles, polydispersity index, and Zeta potential of the PMV were obtained from Dynamic light scattering analysis, using the Zetasizer Nano ZS (Malvern Instruments, UK).

Negative staining TEM technique was used to visualize the vesicles. About 7 μl suspension of vesicles (1mg ml^-1^) was allowed to adsorb onto a freshly glow-discharged carbon-coated grid (400 mesh; Copper), and excess solution was blotted off the grid using Whatman No. 1 filter paper. Negative staining was performed using 2% (w/v) aqueous uranyl acetate, followed by air-drying. The grids were analysed under a Jeol JEM 1400 TEM (Tokyo, Japan) operating at 120 kV. Images were acquired using an Orius SC200B CCD camera and Digital Micrograph software (Gatan, UK).

### 5.4 Estimation of Total Sialic Acid Content of PMVs

The total sialic acid levels of PMVs were estimated using a Sialic acid assay kit (Merck, MAK314-1KT) as per the manufacturer’s instructions.

### 5.5 Extrusion of PMV

Crude plasma membrane vesicles were extruded to generate vesicles of defined sizes. Crude PMVs were diluted to a desired concentration and then passed through polycarbonate membrane filters 23 times with defined pore sizes (400nm and 100nm) (Avanti Polar Lipids, US) using a mini-extruder (Avanti Polar Lipids, US). The extrusion yielded vesicles having a diameter near the expected pore size, and the size was assessed by dynamic light scattering using the Zetasizer Nano ZS (Malvern Instruments, UK).

### 5.6 Cytotoxicity Assay of PMV

To assess the cytotoxicity of PMVs, 1.5 X 10^4^ cells/well (A549 or MDCK or RAW 264.7 cells) were seeded in a 96-well tissue culture plate and allowed to grow as a monolayer for 24 hours. After adherence, the cells were incubated with different concentrations of the plasma membrane vesicles (derived from A549, MDCK, and HEK293T cells) and allowed to grow for 48 hours at 37°C and 5% CO_2_. After 48 hours, cell viability was assessed by MTT assay.

### 5.7 Anti-viral activity of PMV

MDCK and A549 cells were plated in 96-well microtiter plates (10,000 cells/well for A549 and 15,000 cells/well for MDCK). Virus (H3N2 and H1N1) was mixed with varying amounts of PMVs in Serum-free media (SFM) containing TPCK-treated trypsin (0.5 μg ml^-1^) and was incubated for 1 hour at 4°C. To induce infection with the virus, cells were washed with 1XPBS (pH 7.4), then different dilutions of vesicles and the same amount of virus were added to each well and incubated for 45 minutes at 37°C, 5% CO_2_. After removing infection media, cells were washed twice with 1XPBS, and growth media containing 0.1% FBS, 0.3% BSA, and trypsin (2.5 μg ml^-1^) was added, and cells were incubated for 48 hours at 37°C, 5% CO_2_. Next 10μl of MTT (SRL) (from a stock of 5 mg ml^-1^ in 1X PBS) was added to each well, and the plate was incubated for 4 hours at 37 °C, 5% CO_2_. After that, 100 μl of DMSO was added to each well to dissolve the formazan crystals formed in the process, and absorbance was measured at 570 nm. The wells containing only cells and cells infected with the virus were considered as cell control and virus control, respectively. Cell viability was determined by considering that the cell control absorbance corresponds to 100 % viability. The percentage of cytopathic effect (CPE) inhibition was defined as: [(O.D. _vesicle_ - O.D. _virus control_) / (O.D. _cell control_ - O.D. _virus control_)] ×100, where O.D. _vesicle_ is the absorbance in the presence of virus and vesicle, O.D. _virus control_ is the absorbance in the presence of virus only and O.D. _cell control_ is the absorbance of cell control wells. Each O.D. value was background-subtracted (Zhang et al., 2020), (Panda et al., 2025). The reduction in CPE is plotted as a function of vesicle concentration using GraphPad Prism 6 (San Diego, CA, USA).

### 5.8 Western blot Analysis

For western blot analysis, we selected the virion-associated matrix protein M1, the most abundant structural protein in the influenza viral particle, which is detectable shortly after infection (Eierhoff et al., 2010). To determine M1 expression level in infected cells, A549 or MDCK cells were seeded in a 6-well plate at a cell density of 0.25 × 10 ^6^/ well, and infected with Influenza A virus in the absence and presence of varying concentrations of PMV as described above. After 48 hours of infection, cells were observed for cytopathic effect. Media was harvested for HA titer determination. After removing the media, cells were washed with ice-cold PBS and scraped in a lysis buffer (RIPA-HIMEDIA) in the presence of a protease inhibitor cocktail (SIGMA). To prepare the cell lysates, cells were subjected to sonication for 90 seconds (10 pulses, Amplitude 60%) using a SONICS Vibra-Cell probe sonicator (SONICS). The cells were then pelleted by centrifugation at 16,000 RCF for 20 minutes at 4°C. The supernatant was collected and stored at -20°C. Total protein concentration was determined by bicinchoninic acid assay, using a kit (Thermo Scientific, USA). Cell lysates were separated on SDS-PAGE (12%) and then transferred to a PVDF membrane (Bio-Rad). The membrane was blocked by dipping in 3% skimmed milk (Bio-Rad) in PBS containing 0.1% Tween 20 (Amresco) (1xPBST) for 1 hour with slow rocking at room temperature (RT). IAV-associated matrix protein (M1) was detected by an anti-M1 antibody (1:1000) as primary, and an HRP-conjugated rabbit anti-mouse IgG H&L (1:10,000) was used as a secondary antibody. The same blots were used after stripping for estimating β-actin as a loading control using an HRP-conjugated anti-β-actin (A3854) (1:10,000). Quantification of Western blot was done using ImageJ (Rasband, W.S., ImageJ, U. S. National Institutes of Health, Bethesda, Maryland, USA). The band intensity values obtained were normalised to the corresponding β-actin.

### 5.9 Immunofluorescence Assay

A549 cells were plated (0.05 X 10^4^ cells/well) in a 24-well tissue culture plate containing 12 mm sterile coverslips preplaced before seeding. Virus (Influenza-A/H3N2/X-31) with different PMVs made in Serum-free media (SFM) containing TPCK-treated trypsin (0.5 µg ml^-1^) were incubated for 1 hour at 4°C. To infect with the virus, cells were washed with 1X PBS (pH 7.4), then the virus-PMV mixture was added to each well and incubated for 45-50 minutes at 37°C, 5% CO_2_. After removing the infection media, cells were washed twice with 1X PBS. Next, growth media containing 0.1% FBS, 0.3% BSA, and trypsin (2.5 μg ml^-1^) was added and incubated for 15 hours at 37°C, 5% CO_2_. For immunostaining, cells were first fixed using 4% paraformaldehyde, pH 7.4 (SIGMA) made in 1XPBS, for 10 minutes, and then permeabilized for 15 minutes at room temperature with 0.1% Triton X-100 (SRL) in 1XPBS. Cells were washed with 1X PBS, and blocking solution (2% BSA in 1X PBST) was added for 1 hour at RT. A primary mouse monoclonal Anti M1 antibody (ab22396, Abcam) was added in a 1:1000 dilution for 90 minutes at RT. After that, the cells were washed five times with 1X PBS. Following this, cells were incubated with an Alexa Fluor®488 conjugated secondary antibody (ab150113 from Abcam) diluted in 1% BSA in PBST (1:10000) for 1 hour in the dark. Cells were washed five times with PBST and mounted on glass slides using mounting media (fluoroShield with DAPI, SIGMA). The confocal images were acquired using a confocal laser-scanning microscope (OLYMPUS FV 1200, Japan). Post-acquisition, the images were adjusted for brightness and contrast to enhance clarity and the optimal visualization of fluorescent signals, and were applied uniformly across all samples.

### 5.10 Virus Internalization Assay

To monitor influenza A virus entry, MDCK/A549 cells were seeded at a density of 0.25 ×10[cells per well in 6-well plates. After allowing the cells to adhere overnight, they were infected with influenza A virus in the presence or absence of varying concentrations of vesicles, as previously described. The infection was carried out at 37°C with 5% CO[for 45 minutes to facilitate viral entry. Post-infection, the viral inoculum was removed. De novo protein synthesis was inhibited by cycloheximide in the infection media. Cells were washed with ice-cold PBS to eliminate residual virus particles. Subsequently, cell lysates were prepared (as described above) for western blot analysis. For immunostaining, after 45 minutes of infection under the same conditions, cells were fixed with 4% paraformaldehyde (pH 7.4) for 10 minutes at room temperature. Following fixation, staining was done as per the protocol described earlier.

### 5.11 Hemagglutination Assay

The hemagglutination assay quantifies the strength of the virus in terms of HA titer (Wu et al. 2016). A hemagglutination assay was performed with chicken RBCs, which were isolated by diluting freshly drawn chicken blood in PBS, followed by pelleting at 2500 RCF for 10 min at 25°C and repeated 3-4 times until a clear supernatant was obtained. The supernatant was removed, and RBCs were stored at 4°C. For the HA assay, 50µl of 1X PBS was added from the 2nd to 12th well of a 96-well U-bottom plate. 100µl of the infected media (in the presence and absence of the vesicles) was pipetted into the first well and 2-fold serially diluted across the row. 50 µl of infected media was removed from the last well. Afterward, 50µl of standardized RBC (5% of RBCs in PBS) was mixed in each well and incubated for 30 minutes at RT. The endpoint of agglutination was observed and recorded (Wu et al., 2016).

### 5.12 Hemagglutination Inhibition Assay

The hemagglutination inhibition (HI) assay quantifies the virus binding to the vesicles, resulting in hemagglutination inhibition. In this assay, PMVs were two-fold serially diluted, and an equal amount of virus was added to each well and incubated for 30 minutes at 4°C. 50 µl of 5% RBCs was added to each well and left for 30 minutes at RT. In the absence of PMVs, RBCs hemagglutinate by the influenza virus and do not precipitate at the bottom of the well. However, in the presence of the vesicles, influenza viral particles will no longer be available to agglutinate RBCs, and therefore, RBCs precipitate at the bottom of the well and appear as a red dot. The endpoint vesicle dilution till which point RBCs precipitate was recorded as the HI titer.

## Acknowledgements

This work was supported by the DBT Ramalingaswami Re-entry Fellowship (BT/RLF/Re-entry/11/2020) to S.H., B.Q. and V.K. thank the CSIR and ICMR for the Research Fellowships, respectively. S.H. and R.K.T. thank the Division of Virus Research and Therapeutics, CSIR-CDRI, for providing the infrastructure. X-31 Influenza A virus was a kind gift from Dr. Joshua Zimmerberg from NIH, Bethesda. WSN (H1N1) virus was a kind gift from Dr. Arindam Mondal, IIT Kharagpur. We thank the Sophisticated Analytical Instrument Facility (SAIF), CSIR-CDRI, for providing us with the Confocal Microscopy (Intravital) facility. S.H. thanks the Manipal Institute of Virology for providing the necessary infrastructure.

## Author contribution

BQ designed and performed experiments, analyzed data, generated figures and wrote the first draft; VV performed experiments and generated figures, VK performed vesicle extrusion experiments, GP performed TEM experiments under the supervision of KM, RKT provided resources and supervision, SH conceptualized, designed and performed experiments, analyzed data and generated figures, acquired funding, supervised the overall project, and wrote the complete manuscript with inputs from all the authors.

## Data availability statement

All the data reported are available in the main text and supplementary figures.

## Supplementary Materials

**Figure S1:**
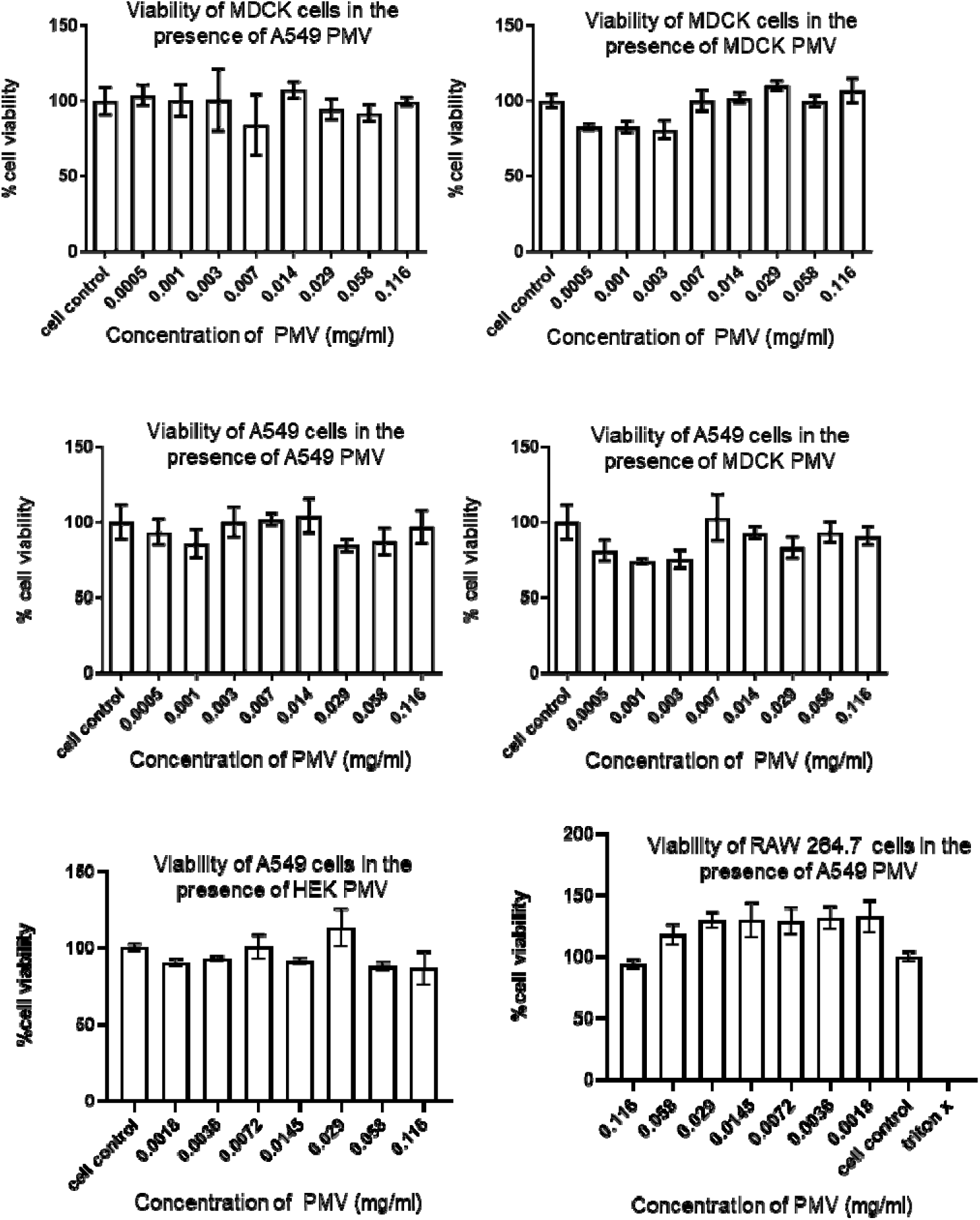
Cytotoxicity of PMV derived from A549, MDCK, and HEK cells in A549, MDCK, and RAW 264.7 cells after 48 hours of co-incubation. Cell viability was measured using an MTT assay, normalized to the control, and reported as mean ± SEM from at least 3 replicates. See the material and method section for details.

**Figure S2.**
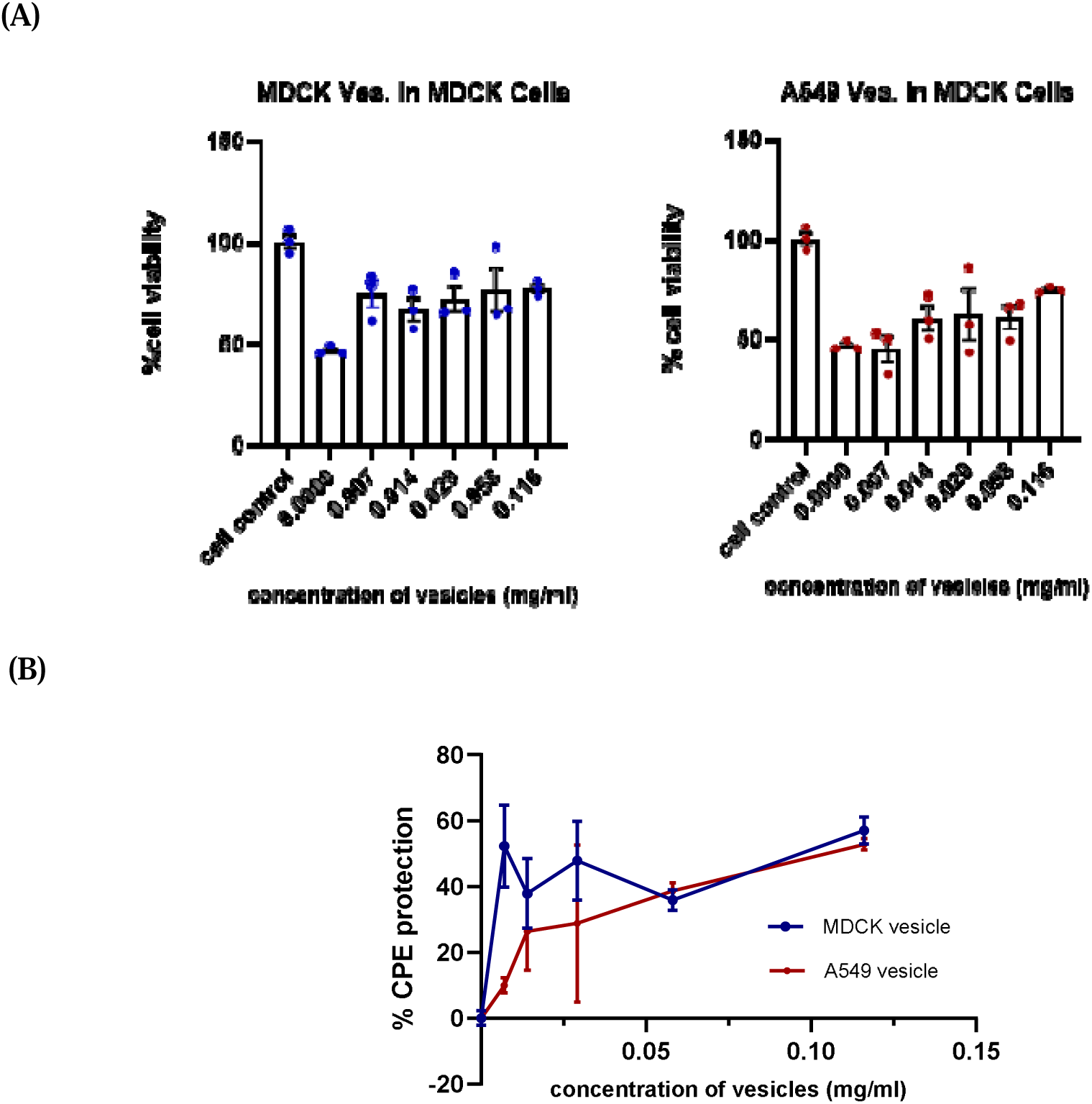
Quantification of *in-vitro* anti-influenza activity of PMV in MDCK cells. (A) Protection from X-31/H3N2 virus-induced CPE (increase in cell viability) by PMV derived from A549, and MDCK cells. (B). Dose-dependent reduction in X-31 infection in MDCK cells, as measured by protection from CPE, as a function of PMV concentration. All other conditions are same as Figure 4.

**Figure S3.**
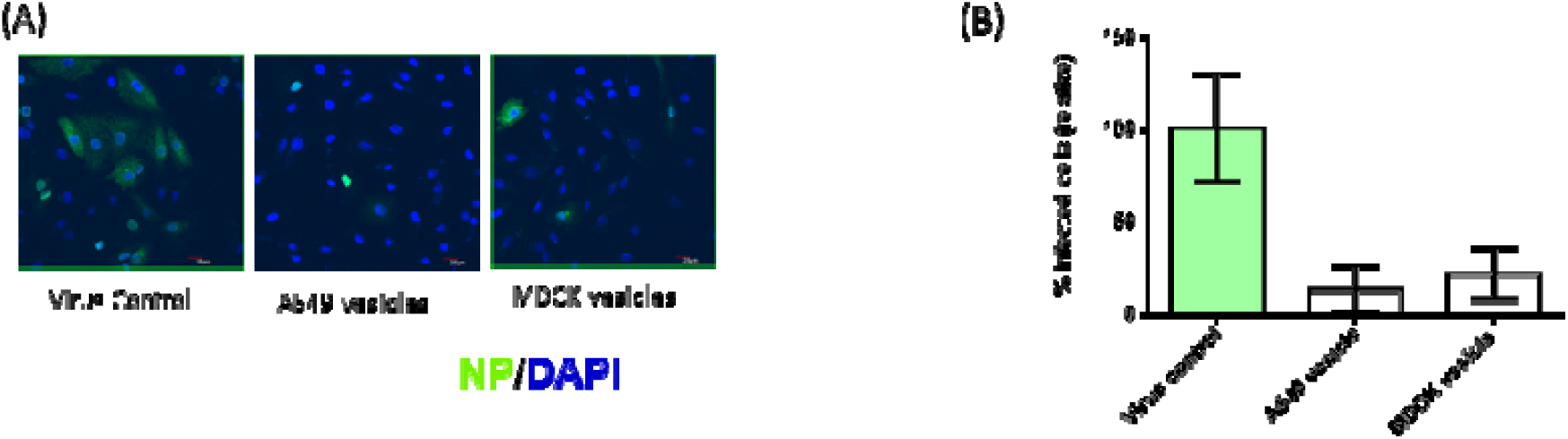
PMV-induced reduction in Influenza A viral (X-31/H_3_N_2_) infection in A549 cells. Infected cells were identified by confocal imaging of the Influenza N protein (green) while cells were marked by DAPI (blue). **(A)** Confocal microscopic images show a decrease in N protein viral protein expression; **(B)** Percentage of infected cells in the presence of PMVs based on N protein staining. (Data represent mean ± SEM of different fields).

**Figure S4.**
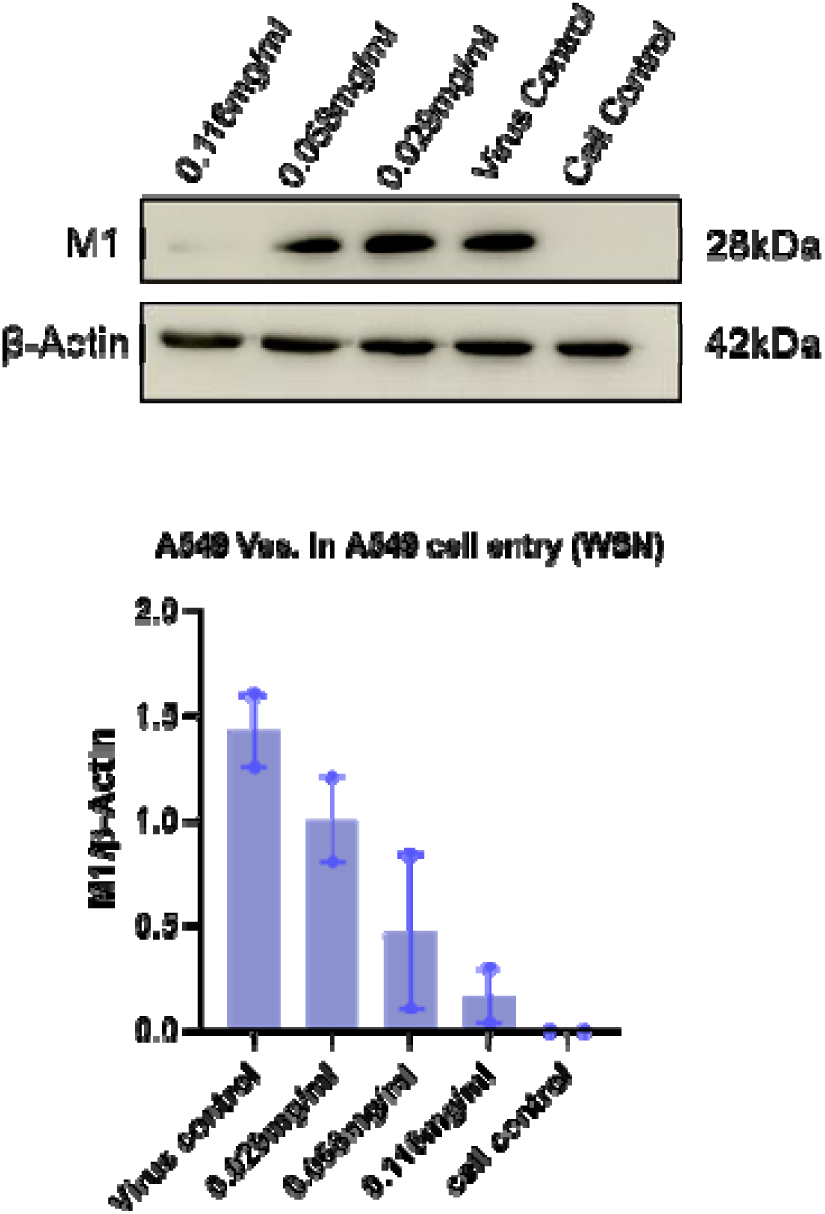
PMV restricts viral entry: A representative western blot showing dose-dependent reduction in influenza A M1 level (45 minutes post-infection) in A549 cells in the presence of PMV derived from A549 cells. β-actin was taken as a loading control. Quantification of M1 levels in A549 cells in the presence of A549 vesicles. See Figure 7 and the materials and methods section for other details.

**Figure S5.**
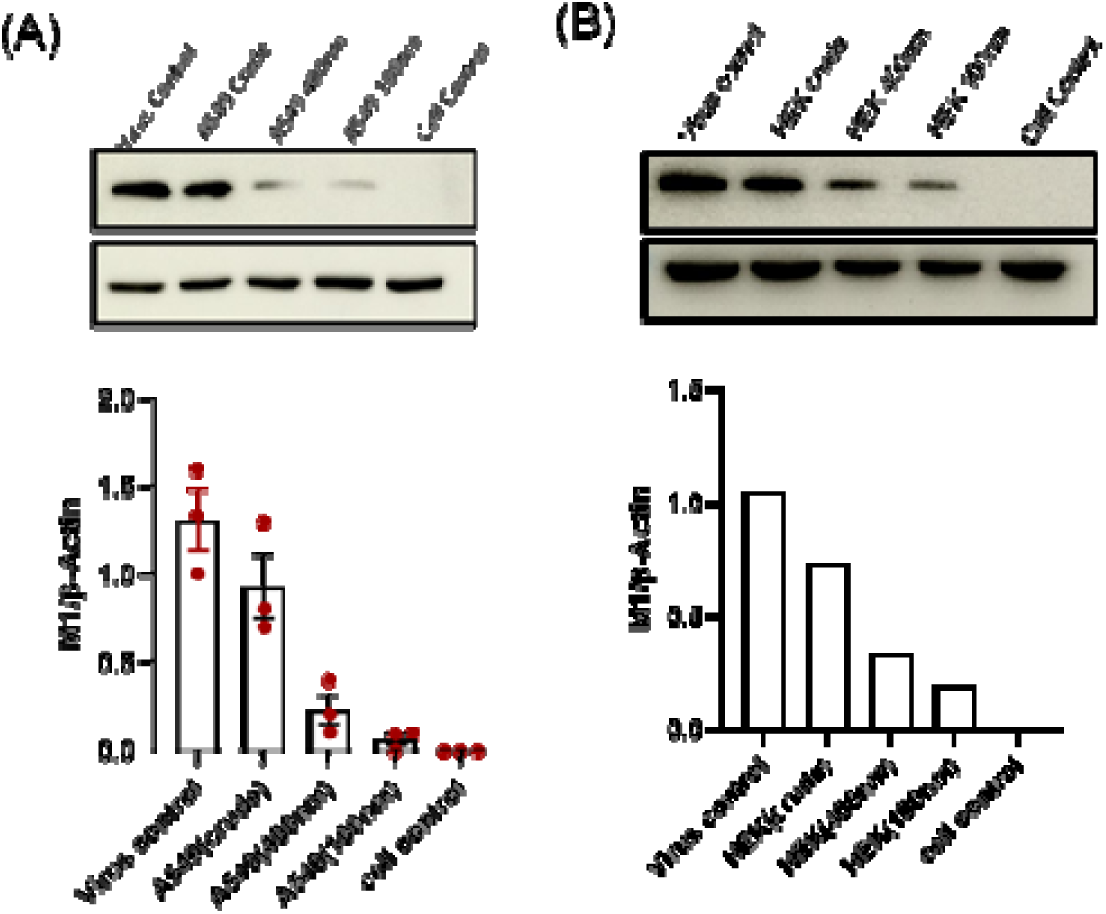
PMV restricts viral entry in a size-dependent manner. Reduction in influenza A M1 level (45 minutes post-infection) in A549 cells in the presence of PMV derived from A549 and HEK cells, shown by western blotting. Representative western blots and corresponding quantification of Influenza A/ WSN/ H1N1 M1 level in the presence of (A) A549 and (B) HEK PMV. β-actin was taken as a loading control. Quantification data represent mean ± SEM of three independent experiments. All other conditions are the same as Figure 8(C).

## References

Alter CL, Detampel P, Schefer RB, et al (2023) High efficiency preparation of monodisperse plasma membrane derived extracellular vesicles for therapeutic applications. Commun Biol 6:478. 10.1038/s42003-023-04859-2

Bai R-H, Lin C-C, Lin C-W (2025) Enzymatic Reactions Dictated by the 2D Membrane Environment. J Phys Chem Lett 16:6745–6756. 10.1021/acs.jpclett.5c00988

Banerjee I, Miyake Y, Philip Nobs S, et al (2014) Influenza A virus uses the aggresome processing machinery for host cell entry. Science 346:473–477. 10.1126/science.1257037

Bhatia S, Camacho LC, Haag R (2016) Pathogen Inhibition by Multivalent Ligand Architectures. J Am Chem Soc 138:8654–8666. 10.1021/jacs.5b12950

Bhatia S, Hilsch M, Cuellar-Camacho JL, et al (2020) Adaptive Flexible Sialylated Nanogels as Highly Potent Influenza A Virus Inhibitors. Angew Chem Int Ed 59:12417–12422. 10.1002/anie.202006145

Bhatia S, Lauster D, Bardua M, et al (2017) Linear polysialoside outperforms dendritic analogs for inhibition of influenza virus infection in vitro and in vivo. Biomaterials 138:22–34. 10.1016/j.biomaterials.2017.05.028

Boni MF (2008) Vaccination and antigenic drift in influenza. Vaccine 26:C8–C14. 10.1016/j.vaccine.2008.04.011

Cail RC, Drubin DG (2023) Membrane curvature as a signal to ensure robustness of diverse cellular processes. Trends Cell Biol 33:427–441. 10.1016/j.tcb.2022.09.004

Carlescu I, Scutaru D, Popa M, Uglea C V. (2009) Synthetic sialic-acid-containing polyvalent antiviral inhibitors. Med Chem Res 18:477–494. 10.1007/s00044-008-9139-7

Costa L, Vilas P, Marcelo B, et al (2019) Antiviral peptides as promising therapeutic drugs. Cell Mol Life Sci 76:3525–3542. 10.1007/s00018-019-03138-w

de Courville C, Cadarette SM, Wissinger E, Alvarez FP (2022) The economic burden of influenza among adults aged 18 to 64: A systematic literature review. Influenza Other Respir Viruses 16:376–385. 10.1111/irv.12963

Eierhoff T, Hrincius ER, Rescher U, et al (2010) The epidermal growth factor receptor (EGFR) promotes uptake of influenza a viruses (IAV) into host cells. PLoS Pathog 6:. 10.1371/journal.ppat.1001099

Eierhoff T, Ludwig S, Ehrhardt C (2009) The influenza A virus matrix protein as a marker to monitor initial virus internalisation. Biol Chem 390:509–515. 10.1515/BC.2009.053

Gray EM, Díaz-Vázquez G, Veatch SL (2015) Growth conditions and cell cycle phase modulate phase transition temperatures in RBL-2H3 derived plasma membrane vesicles. PLoS One 10:1–16. 10.1371/journal.pone.0137741

Haldar S, Okamoto K, Dunning RA, Kasson PM (2020) Precise Triggering and Chemical Control of Single-Virus Fusion within Endosomes. J Virol 95:. 10.1128/JVI.01982-20

Herold S, Becker C, Ridge KM, Budinger GRS (2015) Influenza virus-induced lung injury: Pathogenesis and implications for treatment. Eur Respir J 45:1463–1478. 10.1183/09031936.00186214

Ingato D, Edson JA, Zakharian M, Kwon YJ (2018) Cancer Cell-Derived, Drug-Loaded Nanovesicles Induced by Sulfhydryl-Blocking for Effective and Safe Cancer Therapy. ACS Nano 12:9568–9577. 10.1021/acsnano.8b05377

Irwin KK, Renzette N, Kowalik TF, Jensen JD (2016) Antiviral drug resistance as an adaptive process. Virus Evol 2: vew014. 10.1093/ve/vew014

Lakadamyali M, Rust MJ, Babcock HP, Zhuang X (2003) Visualizing infection of individual influenza viruses. Proc Natl Acad Sci U S A 100:9280–9285. 10.1073/pnas.0832269100

Lampejo T (2020) Influenza and antiviral resistance: an overview. Eur J Clin Microbiol Infect Dis 39:1201–1208. 10.1007/s10096-020-03840-9

Le Q-V, Lee J, Lee H, et al (2021) Cell membrane-derived vesicles for delivery of therapeutic agents. Acta Pharm Sin B 11:2096–2113. 10.1016/j.apsb.2021.01.020

Liu C, Hu L, Dong G, et al (2023) Emerging drug design strategies in anti-influenza drug discovery. Acta Pharm Sin B 13:4715–4732. 10.1016/j.apsb.2023.08.010

Longmire M, Choyke PL, Kobayashi H (2008) Clearance Properties of Nano-Sized Particles and Molecules as Imaging Agents: Considerations and Caveats. Nanomedicine 3:703–717. 10.2217/17435889.3.5.703

Matrosovich M, Klenk H (2003) Natural and synthetic sialic acid-containing inhibitors of influenza virus receptor binding. Rev Med Virol 13:85–97. 10.1002/rmv.372

Matz HC, Ellebedy AH (2025) Vaccination against influenza viruses annually: Renewing or narrowing the protective shield? J Exp Med 222:. 10.1084/jem.20241283

Morens DM, Fauci AS (2020) Emerging Pandemic Diseases: How We Got to COVID-19. Cell 182:1077–1092. 10.1016/j.cell.2020.08.021

Nagao M, Matsubara T, Hoshino Y, et al (2019) Synthesis of Various Glycopolymers Bearing Sialyllactose and the Effect of Their Molecular Mobility on Interaction with the Influenza Virus. Biomacromolecules 20:2763–2769. 10.1021/acs.biomac.9b00515

Ng CY, Kee LT, Al-Masawa ME, et al (2022) Scalable Production of Extracellular Vesicles and Its Therapeutic Values: A Review. Int J Mol Sci 23:7986. 10.3390/ijms23147986

Nogrady B (2025) Virology’s most wanted: the influenza virus. Nature. 10.1038/d41586-025-03607-2

Ogata M, Hidari KIPJ, Kozaki W, et al (2009) Molecular Design of Spacer-N-Linked Sialoglycopolypeptide as Polymeric Inhibitors Against Influenza Virus Infection. Biomacromolecules 10:1894–1903. 10.1021/bm900300j

Panda MS, Qazi B, Vishwakarma V, et al (2025) Developing peptide-based fusion inhibitors as an antiviral strategy utilizing coronin 1 as a template. RSC Med Chem 16:125–136. 10.1039/D4MD00523F

Parshad B, Schlecht MN, Baumgardt M, et al (2023) Dual-Action Heteromultivalent Glycopolymers Stringently Block and Arrest Influenza A Virus Infection In Vitro and Ex Vivo. Nano Lett 23:4844–4853. 10.1021/acs.nanolett.3c00408

Pedersen JC (2014) Hemagglutination-Inhibition Assay for Influenza Virus Subtype Identification and the Detection and Quantitation of Serum Antibodies to Influenza Virus. Methods Mol Biol 1161: 11–25

Purohit A, Chhatre N, Qazi B, et al (2025) Pseudo-Infection Model: A Bottom-Up Synthetic Approach to Cellular Entry of Enveloped Viruses and Virus Mimics. Small 21:. 10.1002/smll.202412218

Putri WCWS, Muscatello DJ, Stockwell MS, Newall AT (2018) Economic burden of seasonal influenza in the United States. Vaccine 36:3960–3966. 10.1016/j.vaccine.2018.05.057

Sempere Borau M, Stertz S (2021) Entry of influenza A virus into host cells — recent progress and remaining challenges. Curr. Opin. Virol. 48:23–29

Sezgin E, Kaiser HJ, Baumgart T, et al (2012) Elucidating membrane structure and protein behavior using giant plasma membrane vesicles. Nat Protoc 7:1042–1051. 10.1038/nprot.2012.059

Sigal GB, Mammen M, Dahmann G, Whitesides GM (1996) Polyacrylamides Bearing Pendant α-Sialoside Groups Strongly Inhibit Agglutination of Erythrocytes by Influenza Virus: The Strong Inhibition Reflects Enhanced Binding through Cooperative Polyvalent Interactions. J Am Chem Soc 118:3789–3800. 10.1021/ja953729u

Skehel JJ, Wiley DC (2000) Receptor Binding and Membrane Fusion in Virus Entry: The Influenza Hemagglutinin. Annu Rev Biochem 69:531–569. 10.1146/annurev.biochem.69.1.531

Spaltenstein A, Whitesides GM (1991) Polyacrylamides bearing pendant.alpha.-sialoside groups strongly inhibit agglutination of erythrocytes by influenza virus. J Am Chem Soc 113:686–687. 10.1021/ja00002a053

von Itzstein M (2007) The war against influenza: discovery and development of sialidase inhibitors. Nat Rev Drug Discov 6:967–974. 10.1038/nrd2400

Wu Y, Cho M, Shore D, et al (2016) Hemagglutination Inhibition (HI) Assay of Influenza Viruses with Monoclonal Antibodies. Bio Protoc 6:1–6. 10.21769/bioprotoc.1828

Zhang P, Li C, Ma X, et al (2024) Glycopolymer with Sulfated Fucose and 6[1-Sialyllactose as a Dual-Targeted Inhibitor on Resistant Influenza A Virus Strains. ACS Macro Lett 13:874–881. 10.1021/acsmacrolett.4c00221

Zhang X, Tu S, Tian J, et al (2025) Liposome-Based Nanoparticle Delivery Systems for Lung Diseases: Opportunities and Challenges. Int J Nanomedicine Volume 20:12485–12509. 10.2147/IJN.S538382

Zhang ZR, Zhang YN, Li XD, et al (2020) A cell-based large-scale screening of natural compounds for inhibitors of SARS-CoV-2. Signal Transduct Target Ther 5:3–5. 10.1038/s41392-020-00343-z

